# Differential antigenic imprinting effects between influenza H1N1 hemagglutinin and neuraminidase in a mouse model

**DOI:** 10.1101/2024.04.17.589918

**Authors:** Huibin Lv, Qi Wen Teo, Chang-Chun D. Lee, Weiwen Liang, Danbi Choi, Kevin J. Mao, Roberto Bruzzone, Ian A. Wilson, Nicholas C. Wu, Chris K. P. Mok

**Affiliations:** Carl R. Woese Institute for Genomic Biology, University of Illinois at Urbana-Champaign, Urbana, IL 61801, USA; Department of Biochemistry, University of Illinois at Urbana-Champaign, Urbana, IL 61801, USA; HKU-Pasteur Research Pole, School of Public Health, Li Ka Shing Faculty of Medicine, The University of Hong Kong, Hong Kong SAR, China; Department of Integrative Structural and Computational Biology, The Scripps Research Institute, La Jolla, CA 92037, USA; Istituto Pasteur Italia, Rome, Italy; Centre for Immunology & Infection, Hong Kong Science Park, Hong Kong SAR, China; Skaggs Institute for Chemical Biology, The Scripps Research Institute, La Jolla, CA 92037, USA; Center for Biophysics and Quantitative Biology, University of Illinois at Urbana-Champaign, Urbana, IL 61801, USA; Carle Illinois College of Medicine, University of Illinois at Urbana-Champaign, Urbana, IL 61801, USA; The Jockey Club School of Public Health and Primary Care, The Chinese University of Hong Kong, Hong Kong SAR, China; Li Ka Shing Institute of Health Sciences, Faculty of Medicine, The Chinese University of Hong Kong, Hong Kong SAR, China; S.H. Ho Research Centre for Infectious Diseases, Chinese University of Hong Kong, Hong Kong, China

## Abstract

Understanding how immune history influences influenza immunity is essential for developing effective vaccines and therapeutic strategies. This study investigates the antigenic imprinting of influenza hemagglutinin (HA) and neuraminidase (NA) using a mouse model with sequential infection by four seasonal H1N1 strains. Our findings reveal that, among pre-2009 H1N1 strains, the extent of infection history correlates with the restriction of antibody responses to antigenically drifted HA, but not NA. This suggests the mouse model failed to recapitulate NA imprinting in humans, likely due to the difference in NA immunodominance hierarchy between humans and mice. Nevertheless, pre-existing antibodies induced by infection with pre-2009 influenza virus impeded both functional HA and NA antibody responses against a 2009 pandemic H1N1 strain. Overall, this study provides insights into antigenic imprinting for influenza virus, as well as the limitations of using mouse models for studying antigenic imprinting.

**Importance:** Influenza viruses continue to pose a significant threat to human health, with vaccine effectiveness being a persistent concern. One important factor is the individual immune history can influence subsequent antibody responses. While many studies have focused on how pre-existing antibodies influence the induction of anti-HA antibodies after influenza virus infections or vaccinations, the impact on anti-NA antibodies has been less extensively investigated. In this study, using a mouse model, we highlighted within the pre-2009 H1N1 strains, a greater extent of immune history negatively affected anti-HA antibodies but positively influenced anti-NA antibody responses. However, for the 2009 pandemic H1N1 strain, which underwent with antigenic shift, both anti-HA and anti-NA antibody responses have been impeded by the antibodies induced by pre-2009 H1N1 virus infection. These findings have important implications for enhancing our understanding of antigenic imprinting on anti-HA and anti-NA antibody response and for developing more effective vaccination strategies.

## Introduction

During each influenza virus infection, the human immune system produces a polyclonal antibody response targeting the two main surface glycoproteins of influenza virus: hemagglutinin (HA) and neuraminidase (NA). HA, the predominant surface antigen, consists of a globular head domain containing the receptor binding site and a stem domain with the molecular machinery to facilitate cell entry through fusion of the viral and host membranes [1]. In contrast, the NA protein aids virus release by cleaving terminal sialic acids, enabling nascent virus particles to detach from the host cell membrane [2]. For a long time, it was believed that an effective humoral immune response to influenza virus primarily involved antibodies against HA. However, recent studies have shown that anti-NA antibodies can also play a substantial antiviral role, independent of the HA antibody response [3–5].

Almost everyone has been infected by influenza virus since their childhood and experiences reinfection by antigenically drifted strains on a regular basis [6]. The antigenic imprinting theory suggests that immune history can influence the magnitude and quality of antibody responses to a subsequent infection [7, 8]. A striking example is during the 2009 H1N1 “swine flu” pandemic, older individuals appeared to exhibit lower relative mortality rates compared to other age groups, possibly attributed to their exposure during childhood to antigenically similar H1N1 strains originating from the 1918 ‘Spanish flu’ pandemic [9, 10]. However, understanding the impact of age-dependent immune history on the antibody response to the 2009 H1N1 pandemic virus remains largely elusive, primarily due to the complexity of human experiences with infection and vaccination.

Due to the abundance of HA on the influenza virus surface, antigenic imprinting is most often applied to anti-HA antibody responses [11–15]. A noteworthy example is that early childhood infections with either H1N1 or H3N2 influenza viruses confer protection against H5N1 and H7N9 viruses later in life. This is likely due to the generation of anti-HA cross-reactive antibodies targeting shared epitopes across these diverse strains [16]. However, antigenic imprinting on NA is less well characterized [17, 18]. Moreover, the effect of the extent of immune history on both HA and NA simultaneously remains unclear.

In this study, we aim to mimic human conditions in mice by sequentially infecting them with up to four antigenically distinct influenza viruses and challenging them with the 2009 H1N1 pandemic virus. We highlight the extent of immune history can influence the induction of both anti-HA and anti-NA antibodies. Additionally, the binding epitopes targeted by anti-NA antibodies following pre-2009 H1N1 virus infection in the mouse model may differ from those observed in humans. These results suggest that consideration of immune history is crucial for vaccine design. Moreover, caution must be exercised when using mouse models to investigate antigenic imprinting effects in humans.

## Results

### Establishment of a Mouse Model for Sequential Infections with Heterologous Influenza (H1N1) Viruses

To mimic human sequential infections, we selected four pre-2009 H1N1 influenza strains: A/USSR/90/1977 (USSR/77), A/Chile/1/1983 (Chile/83), A/Beijing/262/1995 (Beijing/95), and A/Brisbane/59/2007 (Bris/07). These strains, chosen for their role as vaccine seed strains, are antigenically distinct from each other [19]. To minimize genetic background interference, we integrated their HA and NA genes into the A/Puerto Rico/8/1934 (H1N1) virus backbone using the “6+2” reverse genetic approach [20].

Prior to the sequential infection experiments, we assessed the cross-reactive antigenicity of HA and NA from each virus. Eight-week-old BALB/c mice were infected twice, 21 days apart, with four sets of homologous viruses. Plasma samples were collected 21 days after the second infection (Figure 1A). We performed ELISA and microneutralization assay on each sample to evaluate binding and neutralizing capacities against all four H1N1 viruses. Binding assays revealed cross-reactive binding antibodies to the HA in mice (Figure 1B-E), while H1N1 cross-neutralization was minimal against the three heterologous strains compared to the homologous strain (Figure 1F-I). Notably, strong cross-reactive NA inhibition (NI) was observed via enzyme-linked lectin assay (ELLA) in each group (Figure 1J-1M), supporting the hypothesis that antigenic drift in HA and NA may occur asynchronously [21, 22].

**Figure 1.**
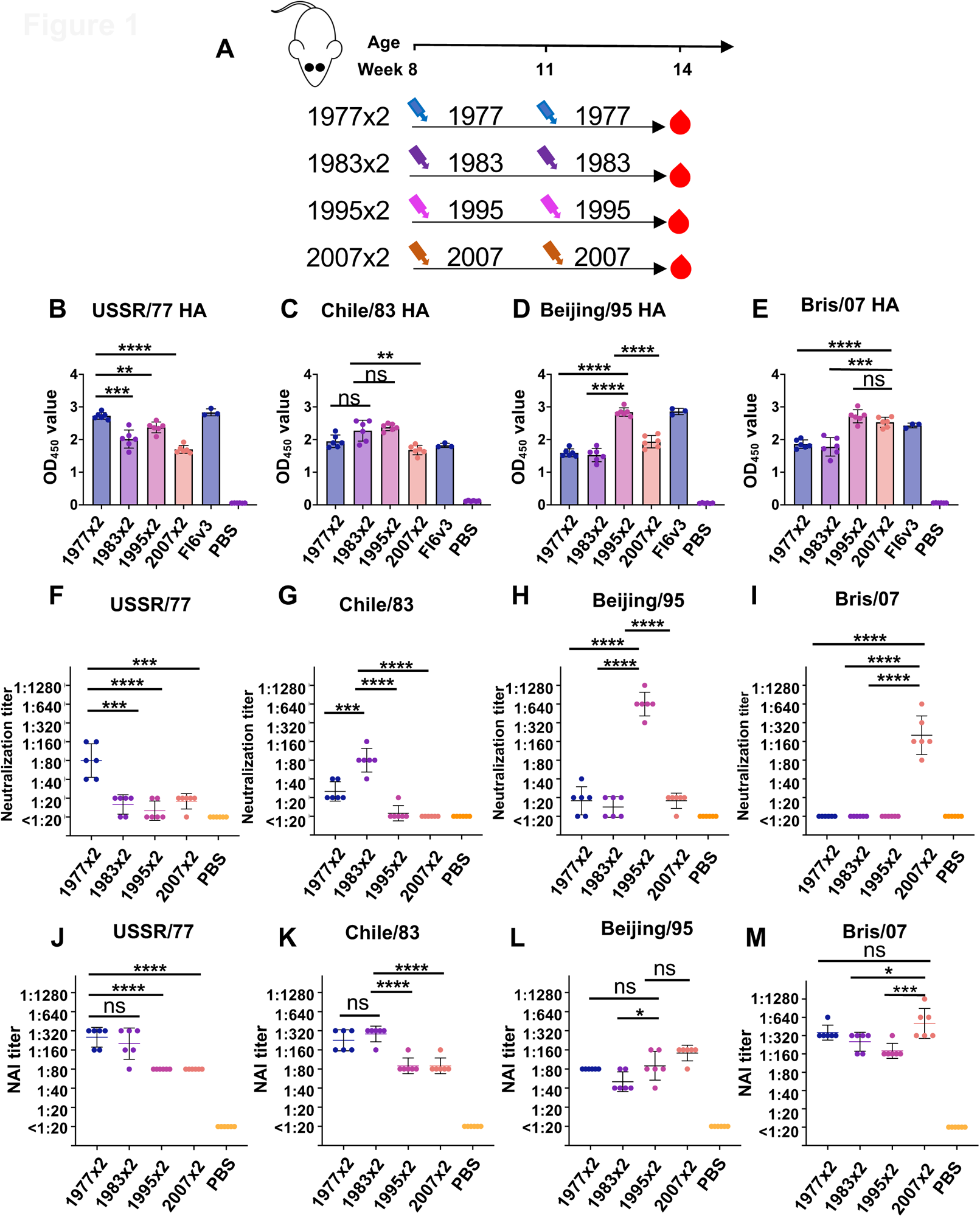
Binding, neutralizing and NAI antibodies induced by sequential homologous viral infection. (A) Experimental design and sample collection. Six mice in each group were inoculated intranasally with sequential homologous H1N1 virus infection strategy (1 × 10^5^ PFU). (B-E) Binding antibodies against (B) USSR/77 HA, (C) Chile/83 HA, (D) Beijing/95 HA, and (E) Bris/07 HA were tested by ELISA. (F-I) Neutralizing antibodies against (F) USSR/77 virus, (G) Chile/83 virus, (H) Beijing/95 virus and (I) Bris/07 virus were assessed by virus neutralization assay. (J-M) NAI antibody against (J) USSR/77 virus, (K) Chile/83 virus, (L) Beijing/95 virus and (M) Bris/07 virus were measured by ELLA. Data are representative of two independent experiments performed in technical duplicate. FI6v3 is an influenza Hemagglutinin (HA) stem-specific antibody, and PBS was used as a negative control. Error bars represent standard deviation. *p*-values were calculated using a two-tailed t-test (\**p* < 0.05, \*\**p* < 0.01, \*\*\**p* < 0.001, \*\*\*\**p* < 0.0001, ns (not significant)).

These findings set the stage for interpreting results from a more comprehensive experimental design involving sequential infection with different heterologous strains. Four-week-old BALB/c mice were divided into four groups: Group 1 was infected once with Bris/07; Group 2 underwent sequential infection with Beijing/95 followed by Bris/07, 12 weeks apart; Group 3 was sequentially infected with Chile/83, Beijing/95, and Bris/07, each 12 weeks apart; Group 4 experienced sequential infection with USSR/77, Chile/83, Beijing/95, and finally Bris/07, again 12 weeks apart (Figure 2A). Two control groups were included: Group 5, infected once with USSR/77 and sampled after 39 weeks; and Group 6, comprising 40-week-old mice infected once with Bris/07. Plasma samples from all groups except Group 5 were collected 21 days post-last infection.

**Figure 2.**
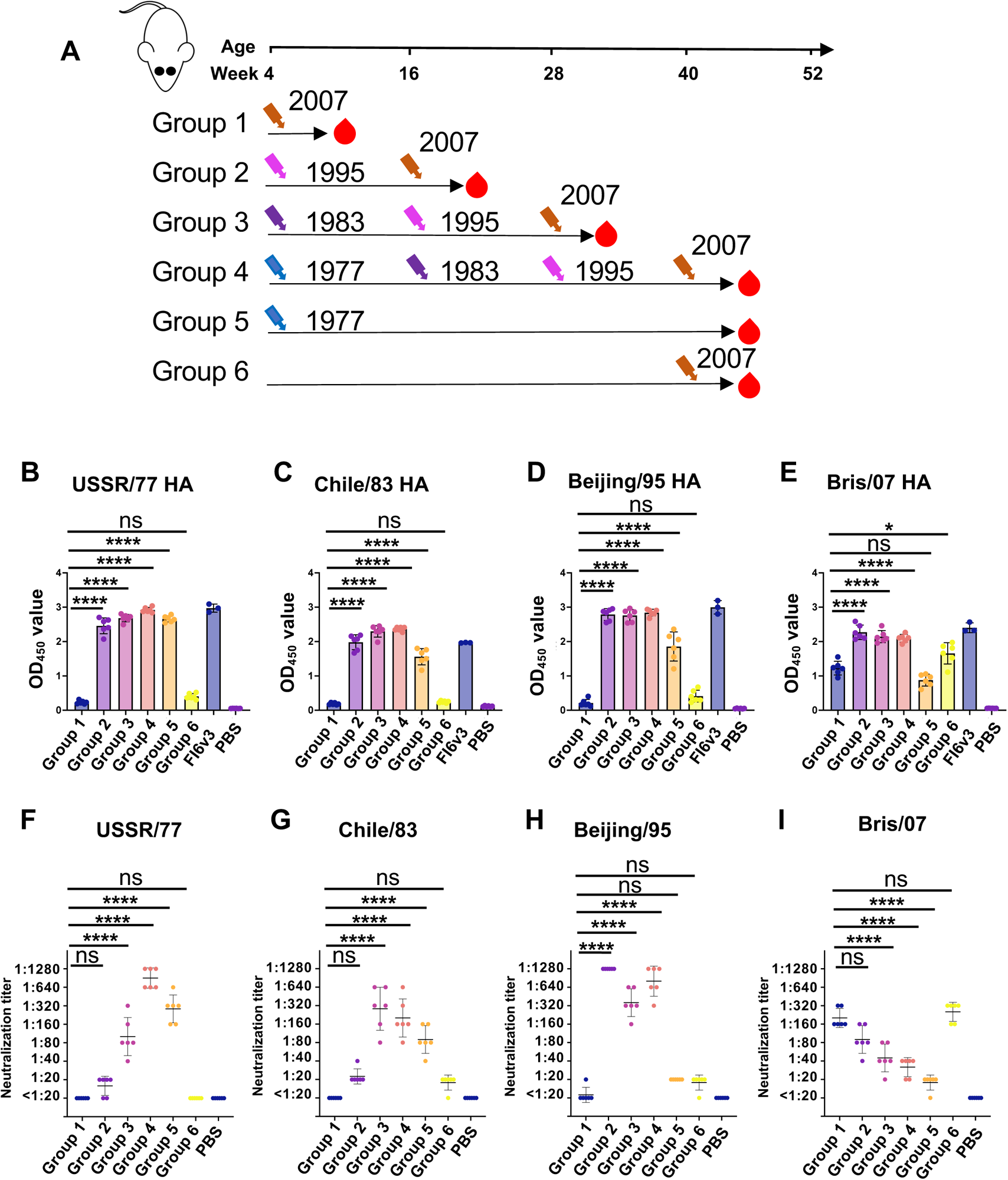
Binding and neutralizing antibodies after sequential viral infection. (A) Experimental design and sample collection. Six mice in each group were inoculated intranasally with sequential H1N1 virus infection strategy (1 × 10^5^ PFU). (B-E) Binding antibodies against (B) USSR/77 HA, (C) Chile/83 HA, (D) Beijing/95 HA, and (E) Bris/07 HA were tested by ELISA. (F-I) Neutralizing antibodies against (F) USSR/77 virus, (G) Chile/83 virus, (H) Beijing/95 virus and (I) Bris/07 virus were assessed by virus neutralization assay. Data are representative of two independent experiments performed in technical duplicate. FI6v3 is an influenza Hemagglutinin (HA) stem specific antibody and PBS was used as a negative control. Error bars represent standard deviation. *p*-values were calculated using a two-tailed t-test (\**p* < 0.05, \*\**p* < 0.01, \*\*\**p* < 0.001, \*\*\*\**p* < 0.0001, ns (not significant)).

### Functional HA and NA Antibodies Show Opposite Trends Following Sequential Infection with Heterologous Influenza Viruses

To investigate antigenic imprinting, we first analyzed plasma binding to the HA proteins of the four viruses. Sequential infection with heterologous H1N1 viruses induced cross-reactive binding antibodies against all four strains (p<0.0001) (Figure 2B-2E). Interestingly, mice infected only with Bris/07 (Group 1) showed lower binding to its cognate HA protein compared to those previously infected with heterologous viruses. Mice in Group 5, infected only with USSR/77, developed cross-reactive binding antibodies to all four viruses (Figure 2B-2E). This suggests that exposure to earlier circulating strains contributes to cross-reactivity to drifted viruses, albeit slightly reduced compared to the parental virus, lasting at least for 43 weeks.

Conversely, neutralizing activity against Bris/07 was highest in mice infected only with this virus (Group 1 and Group 6), and decreased with the number of sequential infections and the distance from the prime to the Bris/07 boost (Figure 2I). This trend suggests a potential relationship between immune priming and viral neutralization activity, where a greater extent of prior infection history may limit the production of neutralizing antibodies. Although Group 5 mice showed relatively strong cross-reactive binding capacity to Beijing/95 and Bris/07 (Figure 2D-2E), no neutralization was observed in the microneutralization assay (Figure 2H-2I), indicating USSR/77 infection-induced antibodies may target non-neutralizing epitopes or the affinity of induced antibodies is relatively low. Comparison of neutralizing antibody responses to Bris/07 in Groups 1 and 6 revealed similarities in immune responses between young and elderly mice (Figure 2I).

Influenza A viruses can be classified into group 1 and 2. We investigated binding cross-reactivity of antibodies towards other human as well as avian group 1 viruses, including A/Puerto Rico/8/1934 (H1N1), A/California/07/2009 (H1N1), A/Japan/305/1957 (H2N2), A/duck/Laos/2006 (H5N1), and A/chicken/Netherlands/2014 (H5N8) (Figure 3B-G). Trends observed were similar to those with the four human H1N1 viruses from 1977 to 2007 (Figure 1B-1E). Using a mini-HA protein derived from the stem domain of Bris/07,[23] we found that stem-binding antibodies may contribute to the targeting of group 1 HAs (Figure 3D). No cross-binding antibody responses were observed against group 2 HA proteins, including those from A/Uruguay/716/2007 (H3N2), A/Anhui/1/2013 (H7N9), and A/Jiangxi/346/2013 (H10N8) (Figure 3H-J), highlighting the specificity of these interactions and the antigenic distinctions within and between these viral groups.

**Figure 3.**
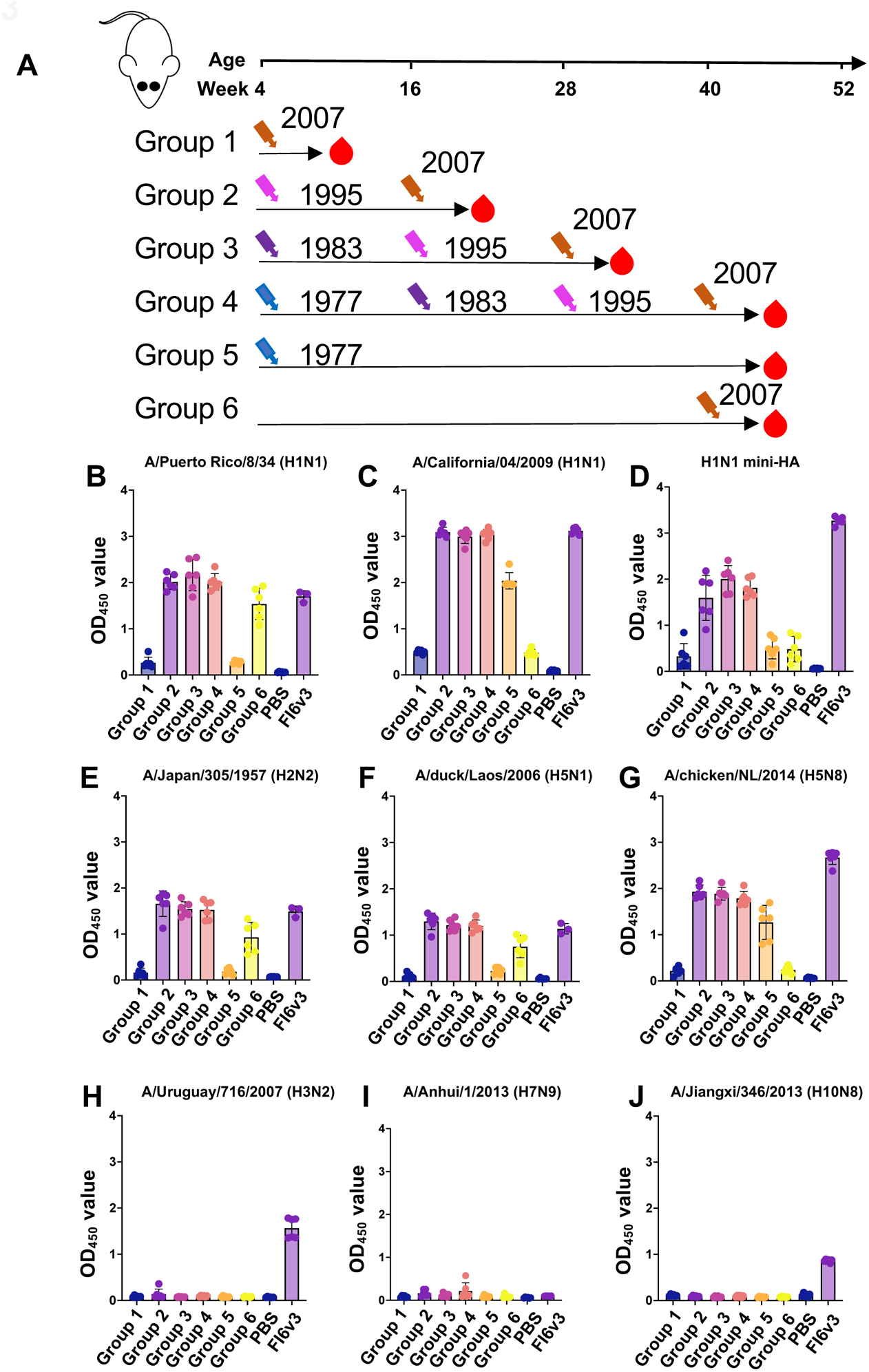
Cross binding antibodies after sequential viral infection. (A) Experimental design and sample collection. Six mice in each group were inoculated intranasally with sequential H1N1 virus infection strategy (1 × 10^5^ PFU). (B-J) Binding antibodies against (B) A/Puerto Rico/8/34 (H1N1) HA, (C) A/California/04/2009 (H1N1) HA, (D) H1N1 mini-HA, (E) A/Japan/305/1957 (H2N2) HA, (F) A/duck/Laos/2006 (H5N1) HA, (G) A/chicken/NL/2014 (H5N8) HA, (H) A/Uruguay/716/2007 (H3N2) HA, (I) A/Anhui/1/2013 (H7N9) HA and (J) A/Jiangxi/346/2013 (H10N8) HA were tested by ELISA. Data are representative of two independent experiments performed in technical duplicate. FI6v3 is an influenza Hemagglutinin (HA) stem specific antibody and PBS was used as a negative control. Error bars represent standard deviation. *p*-values were calculated using a two-tailed t-test (\**p* < 0.05, \*\**p* < 0.01, \*\*\**p* < 0.001, \*\*\*\**p* < 0.0001, ns (not significant)).

Antigenic imprinting is believed to be primarily influenced by HA antibodies, but the role of NA antibodies remains unclear. We analyzed HA and NA via hemagglutination inhibition assay (HAI) and neuraminidase inhibition assay (NAI). Mice with previous heterologous virus infections exhibited lower HAI titers against Bris/07 than with homologous Bris/07 infection (Figure 4A), while a contrasting pattern was observed in NAI results, with heightened functional NAI antibody titers in groups with more infection experiences (Figure 4B). These data suggest more boosts lead to increased antibody responses to conserved sites in NA and show an opposite effect of antigenic imprinting for HA and NA against a specific virus at the same time.

**Figure 4.**
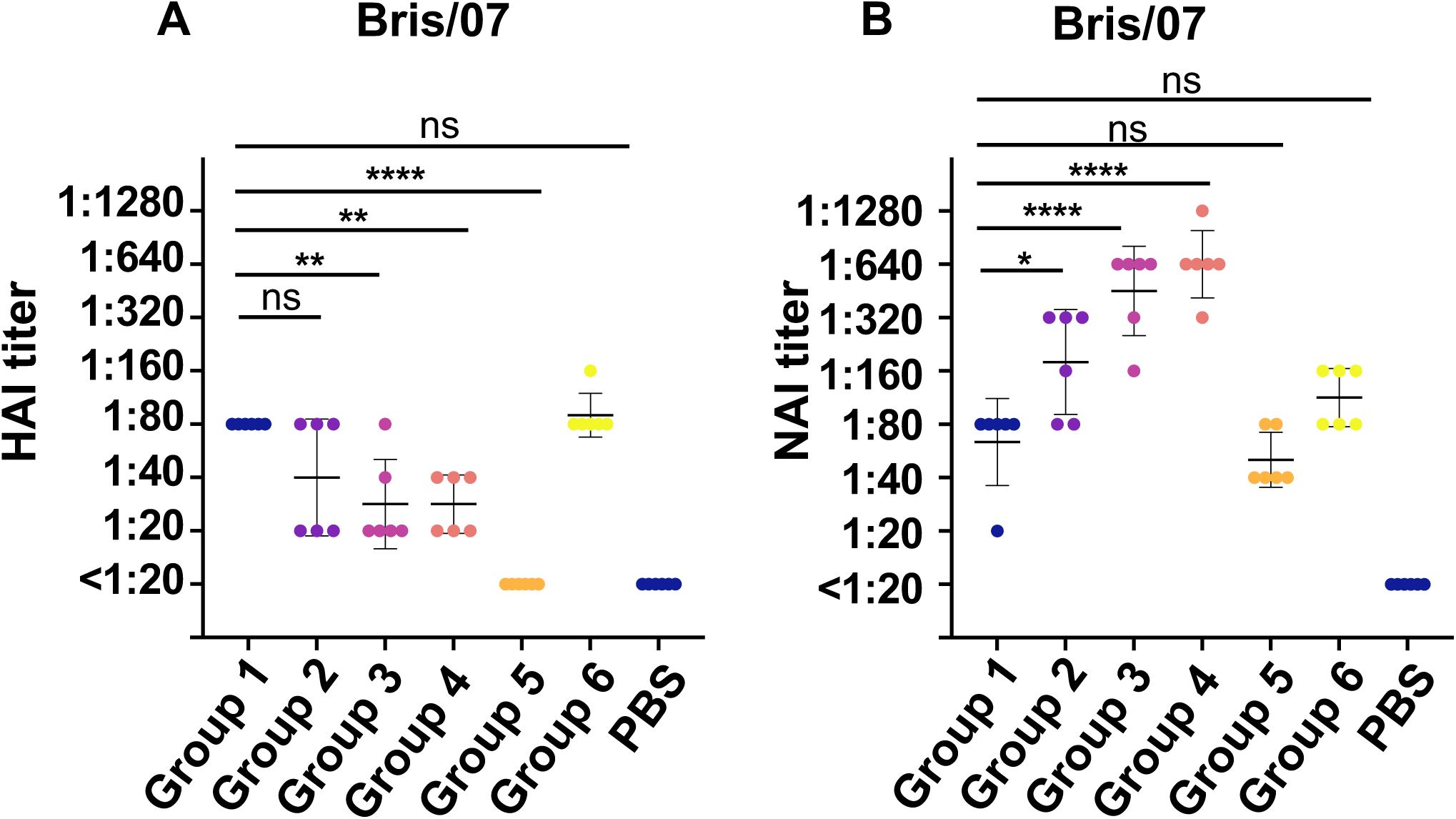
HAI and NAI antibodies after sequential viral infection. (A) Hemagglutination inhibiting antibody against Bris/07 H1N1 virus. (B) Neuraminidase inhibiting antibody against Bris/07 H1N1 virus. Data are representative of two independent experiments performed in technical duplicate. PBS was used as a negative control. Error bars represent standard deviation. *p*-values were calculated using a two-tailed t-test (\**p* < 0.05, \*\**p* < 0.01, \*\*\**p* < 0.001, \*\*\*\**p* < 0.0001, ns (not significant)).

### Impact of Antigenic Shift on Establishment of Antigenic Imprinting

Sequential infection with four strains induced cross-binding antibodies against Cal/09, but no neutralization activity was observed (Figure 3C and S1A). This raises questions about the role of antigenic shift in the development of antigenic imprinting, particularly during the 2009 pandemic. When we challenged mice from Groups 1-4 with a lethal dose of Cal/09 (Figure 5A), all previously infected mice provided 100% protective efficacy in body weight recovery and survival (Figure S2A-S2B). A significant reduction in viral load in the lungs was observed in Group 2-4 which the mice have more than two rounds of heterologous infection (Groups 2-4) (*p*<0.05) (Figure S2C). Plasma collected 21 days post Cal/09 viral infection showed diminished neutralization, HAI, and NAI against Cal/09 for heterologous immunization compared to Bris/07 homologous immunization (Figure 5B-5D), suggesting concurrent antigenic imprinting phenomena induced by shifts in both HA and NA genes. It is interesting that we also observed this imprinting effect is less on NAI, compared to neutralizing activity and HAI (Figure 5D).

**Figure 5.**
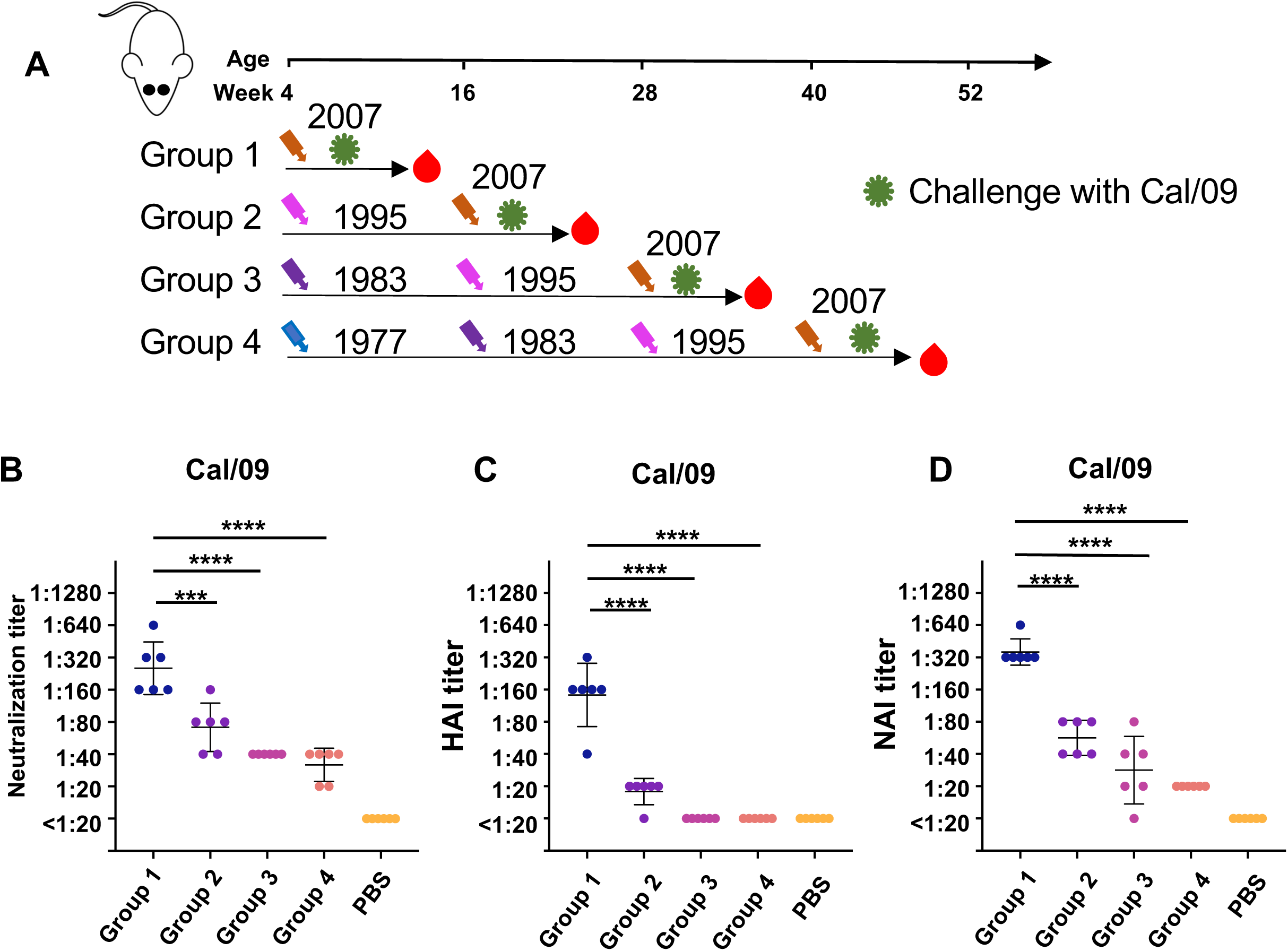
Neutralizing, HAI and NAI antibodies with sequential infection history after Cal/09 H1N1 challenge. (A) Experimental design and sample collection. Six mice in each group were first inoculated intranasally with sequential H1N1 virus infection strategy (1 × 10^5^ PFU) and were challenged with Cal/09 H1N1 virus (4 × 10^5^ PFU). (B) Neutralizing antibodies against Cal/09 H1N1 virus were assessed by virus neutralization assay. (C) Hemagglutination inhibiting antibody against Cal/09 H1N1 virus. (D) Neuraminidase inhibiting antibody against Cal/09 H1N1 virus. Data are representative of two independent experiments performed in technical duplicate. PBS was used as a negative control. Error bars represent standard deviation. *p*-values were calculated using a two-tailed t-test (\**p* < 0.05, \*\**p* < 0.01, \*\*\**p* < 0.001, \*\*\*\**p* < 0.0001, ns (not significant)).

To understand the mechanism of antigenic imprinting against Cal/09 NA, we compared amino acid residues in the NA of Cal/09 with those of the four pre-2009 H1N1 strains. We focused on amino acid residues that are completely conserved across the four pre-2009 NAs of interest, but differed in Cal/09 NA (Figure 6A). These residues are highlighted on the surface of Cal/09 NA structure (Figure 6B). Many of these mutations surround the NA active site, such as I149, N220, Q249, K342, S343, N344 and N372. It is noted that most of these mutations are in the major antigenic sites for the NA protein [24]. Moreover, several studies reported that some of the NA antibodies that bind outside the active site can inhibit NA activity by steric hindrance [3, 25]. On the other hand, the glycosylation profiles have been also changed and may influence the antibody response in Cal/09 NA. For example, NWS at 455-457 in four pre-2009 N1 stains goes to GWS in Cal/09 N1 and 434 where it goes from KTT (1977 and 1983 N1) to NTT (glycan in 1995 and 2007 N1) to NTI (Cal/09). Taken together, NA antibodies induced by sequential infection of pre-2009 viruses in the mouse model may dominantly target to the epitopes located in and around the active site that are conserved in pre-2009 strains, but mutated in Cal/09. Therefore, these imprinted antibodies are escaped by Cal/09 virus. This observation further supports the notion that the antigenic disparity in the NA gene may contribute to antigenic imprinting following infection with the Cal/09 virus.

**Figure 6.**
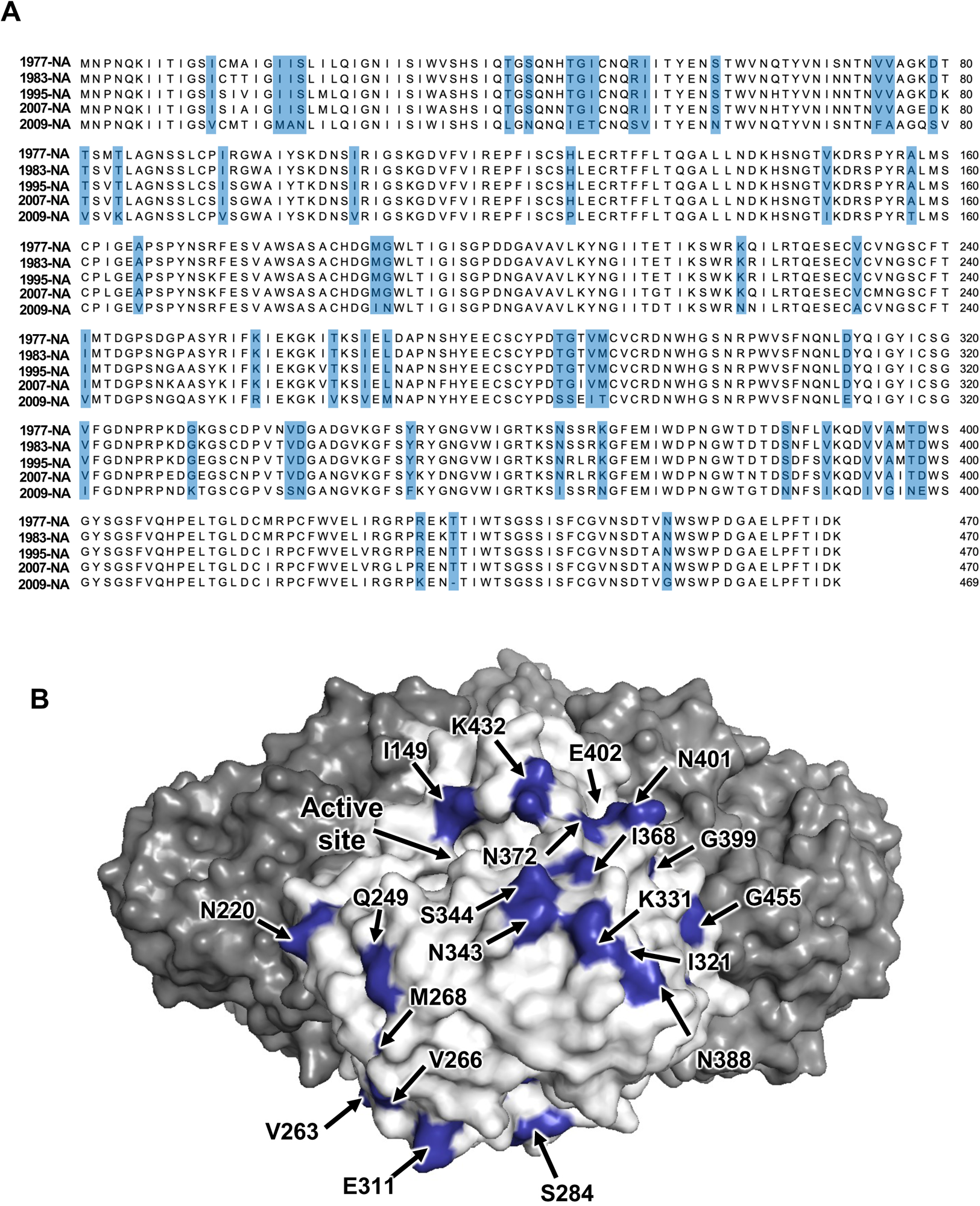
Surface residues difference among pre-2009 H1N1 NA and Cal/09 H1N1 NA. (A) Mutations are highlighted in blue on in a sequence alignment among four pre-2009 N1 protein and Cal/09 NA. (B) Surface residues on Cal/09 NA, which differs from four pre-2009 NAs, are highlighted on the Cal/09 NA protein.

Similar analysis has been performed for pre-2009 and Cal/09 HA amino acid residues (Figure S3A). Residues on the HA head domain are highlighted on the surface of Cal/09 HA structure (Figure S3B). It is interesting that similar types of conserved residues are located closed to the receptor binding site (K145, G158, N159, T187) among four pre-2009 stains, but we don’t observe the same boosting effects in HA after sequential infection, as shown in the NA.

## Discussion

The concept of original antigenic sin (OAS), first described by Thomas Francis Jr in the late 1950s in relation to the influenza virus, has recently been redefined as antigenic imprinting or antigenic seniority [26, 27]. This phenomenon has also been extended to other viruses, such as Dengue virus and SARS-CoV-2 [28–33]. Studies on influenza virus have primarily focused on the HA protein, including monoclonal antibody screening, functional epitope identification, and structural analysis [34–38]. As a result, most observations regarding antigenic imprinting in influenza have focused on the HA. Consequently, the potential impact of antigenic imprinting on the NA protein has been somewhat overlooked.

In our study, we showed antigenic imprinting on the NA protein may produce both positive and negative effects. When dealing with antigenic drift, pre-existing B cell memory from pre-2009 H1N1 viruses may enhance the production of functional NA antibodies against antigenically similar strains through cross-reactivity. However, when encountering a new strain with significant epitope changes due to viral antigenic shift (Cal/09), this pre-existing memory may hinder, but not eliminate, the generation of functional NA antibodies against the new strain. Our findings indicate that pre-existing memory can have dual roles in the context of subsequent viral infections.

Previous research by O’Donnell et al. observed that ferrets with prior seasonal H1N1 infections did not show evidence of original antigenic sin when exposed to the 2009 pandemic H1N1 virus [39]. Their study, focusing on antigenic imprinting, employed a prime-boost strategy but did not explore the influence of the extent of immune history on this phenomenon. In contrast, our study aimed to bridge this gap. We found that a single prior exposure to a pre-2009 H1N1 strain does not influence the neutralizing antibody response to the Cal/09 strain, which is consistent with observations from O’Donnell et al [39]. Additionally, our research sheds light on how infection history affects antibody responses to both HA and NA, emphasizing the need to consider the extent and complexity of the immune history in understanding antigenic imprinting and its implications for influenza virus evolution and immunity.

Another notable aspect of our study is the implication of different immunodominant NA epitopes across various animal species. Daulagala et al. noted lower cross-NAI activity in ferret sera after single H1N1 viral infection with virus strains between 1977 to 1991, while our mouse model displayed apparent cross-reactivity among NA strains from different years [18]. Our hypotheses to explain this observation is the immunodominant epitopes for NA antibody binding induced by ferrets may differ from those induced by mice (Figure 1J-1M). This discrepancy underscores the possibility of species-specific grouping of immunodominant NA epitopes, similar to a pattern also observable in HA. Liu et al. previously demonstrated that, in mice, the antigenic epitopes Sb and Cb2 are immunodominant, while ferret sera predominantly recognize antigenic epitope Sa [40]. Validating the NA immunodominant epitopes and identifying the hierarchy of the NA immunodominant sites in humans could provide valuable information for the rational design of universal vaccines.

In conclusion, our study offers substantial insights into the dynamics of the human immune response to influenza viruses, particularly to both HA and NA. It highlights how the extent of infection history influences antibody responses, a critical factor in the context of antigenic drift and shift. These findings have important implications for enhancing our understanding of influenza and for developing more effective vaccination.

## Materials and methods

### Cells

HEK293T and MDCK cells were cultured in Dulbecco’s Modified Eagle’s Medium (DMEM, high glucose; Gibco) supplemented with 10% heat-inactivated fetal bovine serum (FBS; Gibco), 1% penicillin-streptomycin (Gibco), and 1% Gluta-Max (Gibco). Cells were passaged every 3-4 days using 0.05% Trypsin-EDTA solution (Gibco).

### Protein Expression and Purification

Mini-HA #4900 [23], A/Chile/1/1983 (H1N1) HA, A/Puerto Rico/8/1934 (H1N1) HA, and A/Japan/305/1957 (H2N2) HA proteins were fused with an N-terminal gp67 signal peptide and a C-terminal BirA biotinylation site, thrombin cleavage site, trimerization domain, and Hisx6 tag. These were then cloned into a customized baculovirus transfer vector. Recombinant bacmid DNA was generated using the Bac-to-Bac system (Thermo Fisher Scientific), following the manufacturer’s instructions. Baculovirus was produced by transfecting purified bacmid DNA into adherent Sf9 cells using Cellfectin reagent (Thermo Fisher Scientific), as per the manufacturer’s instructions. The baculovirus was amplified in adherent Sf9 cells at a multiplicity of infection (MOI) of 1. Recombinant proteins were expressed by infecting 1L of suspension Sf9 cells at an MOI of 1. After three days of post-infection, Sf9 cells were centrifuged at 4000 × g for 25 min, and soluble recombinant proteins were purified from the supernatant using Ni Sepharose excel resin (Cytiva), followed by size exclusion chromatography with a HiLoad 16/100 Superdex 200 prep grade column (Cytiva) in 20 mM Tris-HCl pH 8.0, 100 mM NaCl. Proteins were concentrated using an Amicon spin filter (Millipore Sigma) and filtered through 0.22 µm centrifuge Tube Filters (Costar). Protein concentration was determined by Nanodrop (Fisher Scientific), and proteins were aliquoted, flash-frozen in a dry-ice ethanol mixture, and stored at -80°C until use.

HA proteins A/Brisbane/59/2007 (H1N1) (NR-28607), A/California/04/2009 (H1N1) pdm09 (NR-15749), A/duck/Laos/3295/2006 (H5N1) (NR-13509), A/chicken/Netherlands/14015531/2014 (H5N8) (NR-50110), A/Uruguay/716/2007 (H3N2) (NR-15168), A/Anhui/1/2013 (H7N9) (NR-44081), and A/Jiangxi/346/2013 (H10N8) (NR-49440) were obtained from BEI Resources, NIAID, NIH (https://www.beiresources.org/).

### Recombinant Virus Construction and Purification

H1N1 recombinant viruses A/USSR/90/1977 (HA, NA) x A/Puerto Rico/8/1934 (H1N1) (NR-3666), A/Chile/1/1983 (HA, NA) x A/Puerto Rico/8/1934 (H1N1) (NR-3585), A/Beijing/262/1995 (HA, NA) x A/Puerto Rico/8/1934 (H1N1) (NR-3571), and A/Brisbane/59/2007 (HA, NA) x A/Puerto Rico/8/1934 (H1N1) (NR-41797) were obtained from BEI Resources, NIAID, NIH. Recombinant viruses were constructed using a reverse genetics system, as previously described [20]. Briefly, constructed HA and NA DNA plasmids were cloned and transfected into human embryonic kidney 293T cells (ATCC) and Madin-Darby canine kidney (MDCK) cells with a 6-segment plasmid encoding essential viral proteins and virus-like RNA of PR8. Supernatants were injected into 8-10 day old embryonated chicken eggs for viral rescue at 37°C for 48 hours. Viruses were plaque-purified on MDCK cells grown in Dulbecco’s Modified Eagles Medium (DMEM, Gibco) containing 10% fetal bovine serum (FBS, Gibco) and a penicillin-streptomycin mix (100 units/mL penicillin and 100 μg/mL streptomycin, Gibco). Individual plaques were picked, injected into embryonated eggs, and viral RNAs were extracted from allantoic fluids. HA and NA segments were confirmed by Sanger sequencing.

### Mouse Infection and Sample Collection

BALB/c mice were anesthetized with ketamine and xylazine, and intranasally infected with 10^5^ PFU of influenza virus, previously diluted in PBS. Mouse plasma samples were collected in tubes containing heparin as an anticoagulant on day 21 post-infection. The experiments were conducted in the University of Hong Kong’s Biosafety Level 2 (BSL2) facility. The study protocol adhered strictly to the recommendations and was approved by the University of Hong Kong’s Committee on the Use of Live Animals in Teaching and Research (CULATR 5598-20).

### Enzyme-linked immunosorbent assay

Nunc MaxiSorp ELISA plates (Thermo Fisher Scientific) were coated overnight at 4°C with 100 μl of recombinant proteins at 1 μg/mL in 1× PBS. The next day, plates were washed three times with 1× PBS containing 0.05% Tween 20 and blocked with 100 μl of Chonblock blocking/sample dilution ELISA buffer (Chondrex Inc, Redmond, US) for 1 hour at room temperature. Plasma samples, diluted 1:100, were incubated for 2 hours at 37°C. Plates were then washed three times and incubated with horseradish peroxidase (HRP)-conjugated goat anti-mouse IgG antibody (GE Healthcare) diluted 1:5,000 for 1 hour at 37°C. After five washes with PBS containing 0.05% Tween 20, 100 μL of 1-Step™ TMB ELISA Substrate Solution (Thermo Fisher Scientific) was added to each well. Following a 15-minute incubation, the reaction was stopped with 50 μL of 2 M H_2_SO_4_ solution, and absorbance was measured at 450 nm using a Sunrise (Tecan, Männedorf, Switzerland) absorbance microplate reader.

### Microneutralization assay

For the microneutralization (MN) assay, MDCK cells were prepared in each well of 96-well cell culture plates one day before the assay, ensuring a 100% confluent monolayer. Cells were washed once with phosphate-buffered saline (PBS; Gibco) and replaced with minimal essential media (MEM; Gibco) containing 25 mM HEPES (Gibco) and 100 U/mL penicillin-streptomycin (PS; Gibco). All plasma samples for the MN assay were heat-inactivated at 56°C for 30 minutes. Two-fold serial dilutions were performed on the heated plasma to create dilution series ranging from 1:20 to 1:2560. These dilutions were mixed with 100 TCID_50_ of viruses in an equivalent volume and incubated at 37°C for 1 hour. The mixture was then inoculated into cells and incubated at 37°C for another hour. Cell supernatants were discarded and replaced with MEM containing 25 mM HEPES, 100 U/mL PS, and 1 μg/mL TPCK-trypsin (Sigma). Plates were incubated at 37°C for 72 hours, and virus presence was detected by a hemagglutination assay, with results recorded as the MN_50_ titer.

### Hemagglutination-Inhibition (HAI) Assays

Plasma samples were serially diluted two-fold in a 96-well round-bottom plate in a total volume of 25 μl of phosphate-buffered saline (PBS). After dilution, 25 μl of virus [four hemagglutinating units (HAU)] in PBS were added to each well and incubated for 30 minutes. Then, 50 μl of a 1.0% (vol/vol) solution of turkey erythrocytes was added, and the mixture was gently stirred. After 30 minutes at room temperature, the plates were read, and titers were determined as the lowest concentration of monoclonal antibody that fully inhibited agglutination. HAI assays were performed in duplicate.

### Enzyme-linked lectin assay (ELLA)

ELLA experiments were performed as described below. Briefly, each well of a 96-well microtiter plate (Thermo Fisher) was coated with 100 μl of fetuin (Sigma) at a concentration of 25 μg/mL in coating buffer (KPL coating solution; SeraCare) and incubated overnight at 4°C. The following day, 50 μl of plasma samples at the indicated dilution in 2-(N-morpholino) ethanesulfonic acid (MES) buffer (pH 6.5), containing 20 mM CaCl2, 1% bovine serum albumin, and 0.5% Tween 20, were mixed with an equal volume of H1N1 virus. This mixture was added to the fetuin-coated wells and incubated for 18 hours at 37°C. The plate was then washed six times with PBS containing 0.05% Tween 20. Subsequently, 100 μl of horseradish peroxidase-conjugated peanut agglutinin lectin (PNA-HRPO, Sigma–Aldrich) in MES buffer (pH 6.5) with CaCl2 and 1% bovine serum albumin was added to each well and incubated for 2 hours at room temperature in the dark. Following this, the plate was washed six times and developed with 1-Step™ TMB ELISA Substrate Solutions (Thermo Fisher Scientific). The absorbance was measured at 450 nm using a SpectraMax M2 microplate reader (Molecular Devices). Data points were analyzed using Prism software, and the 50% inhibition concentration (IC_50_) was determined as the concentration at which 50% of the neuraminidase (NA) activity was inhibited, compared to the negative control.

## Acknowledgements

This work was supported by Calmette and Yersin scholarship from the Pasteur International Network Association (H.L.), Carl R. Woese Institute for Genomic Biology (IGB) postdoctoral fellpwship (H.L.), Emergency Key Program of Guangzhou Laboratory (EKPG22-30-6) (C.K.P.M), Bill & Melinda Gates Foundation INV-004923 (I.A.W), and NIAID Centers of Excellence for Influenza Research and Response 75N93021C00015 (I.A.W, N.C.W.).

## Author contributions

H.L., R.B., N.C.W. and C.K.P.M. conceived the research idea, planned the study, analysed the data and wrote the manuscript. C.D.L. and I.A.W. provided the purifed HA proteins. H.L., Q.T, and D.C., W.L., K.M., performed the experiments. All authors reviewed and edited the paper.

## Competing Interests

N.C.W. serves as a consultant for HeliXon. The authors declare no competing interests.

---

**Figure S1.**
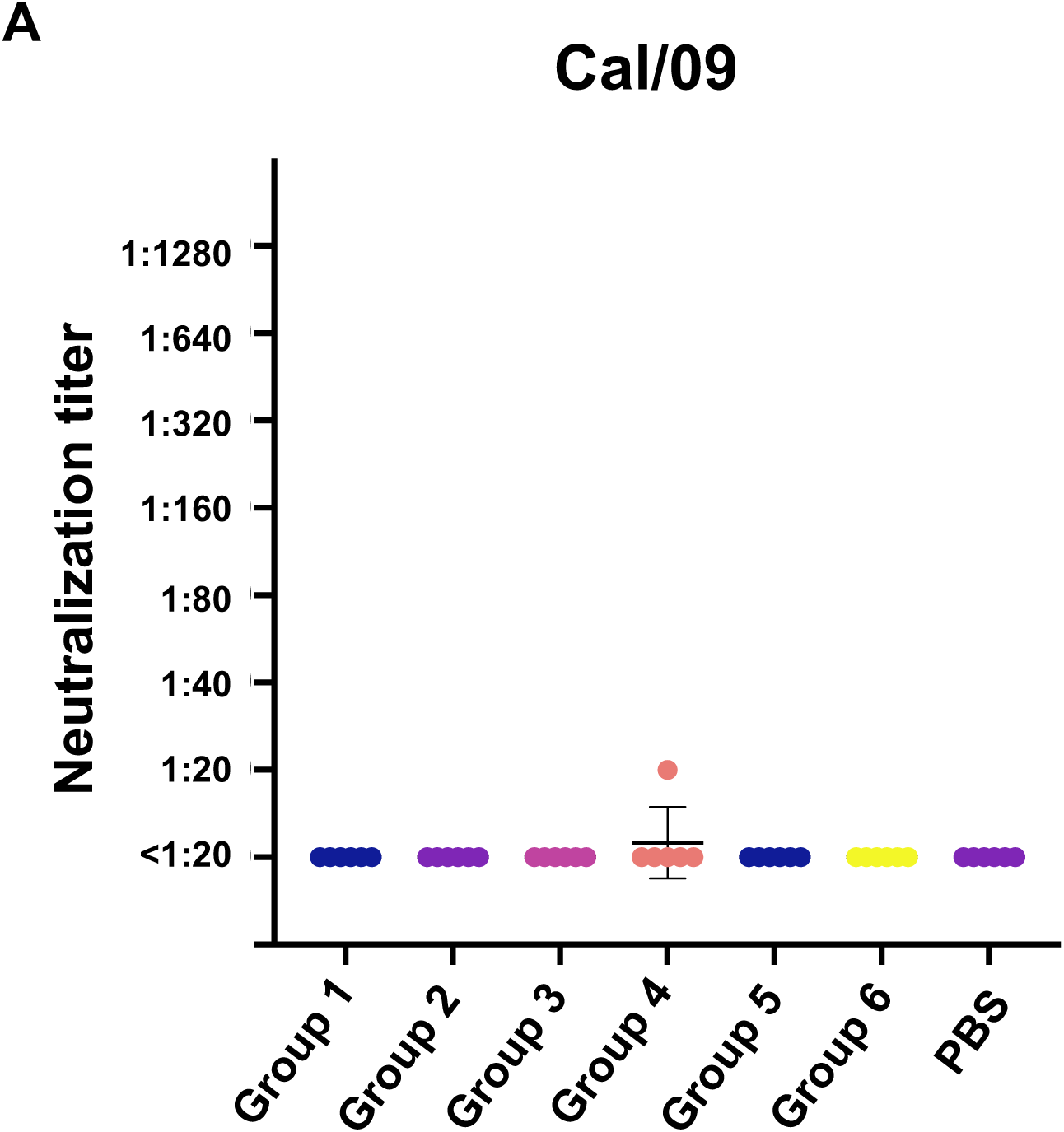
Neutralizing antibodies after sequential viral infection against Cal/09 H1N1. (A) Neutralizing antibodies against Cal/09 virus were assessed by virus neutralization assay. Data are representative of two independent experiments performed in technical duplicate. PBS was used as a negative control.

**Figure S2.**
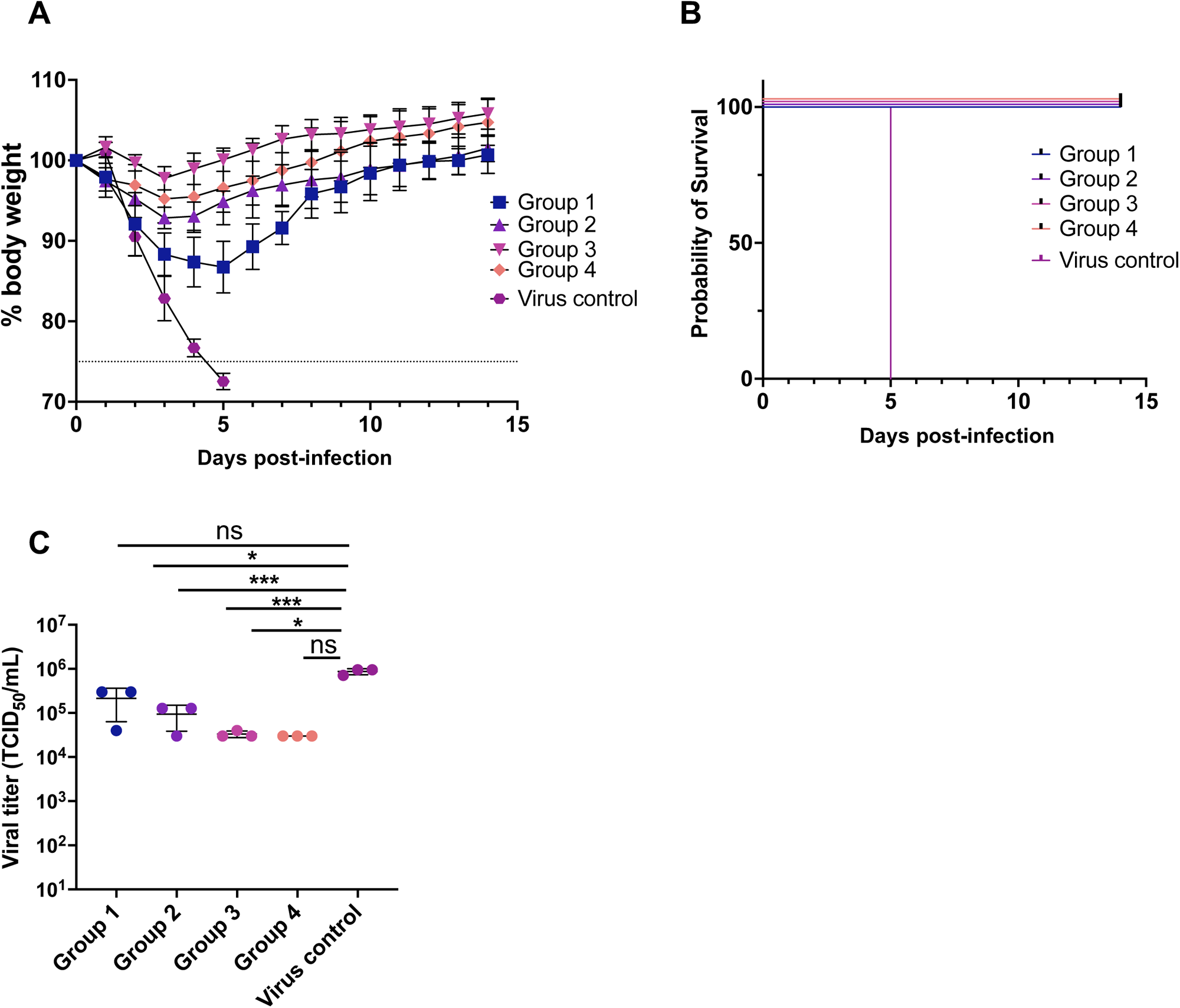
In vivo protection against Cal/09 H1N1 virus after sequential infection. (A) The mean percentage of body weight change post-infection is shown (n = 6). The humane endpoint, which was defined as a weight loss of 25% from initial weight on day 0, is shown as a dotted line. (B) Kaplan-Meier survival curves are shown (n = 6). (C) Lung viral titers on day 3 after infection are shown (n = 3). Solid black lines indicate means ± SD. *p*-values were calculated using a two-tailed t-test (\**p* < 0.05, \*\**p* < 0.01, \*\*\**p* < 0.001, \*\*\*\**p* < 0.0001, ns (not significant)).

**Figure S3.**
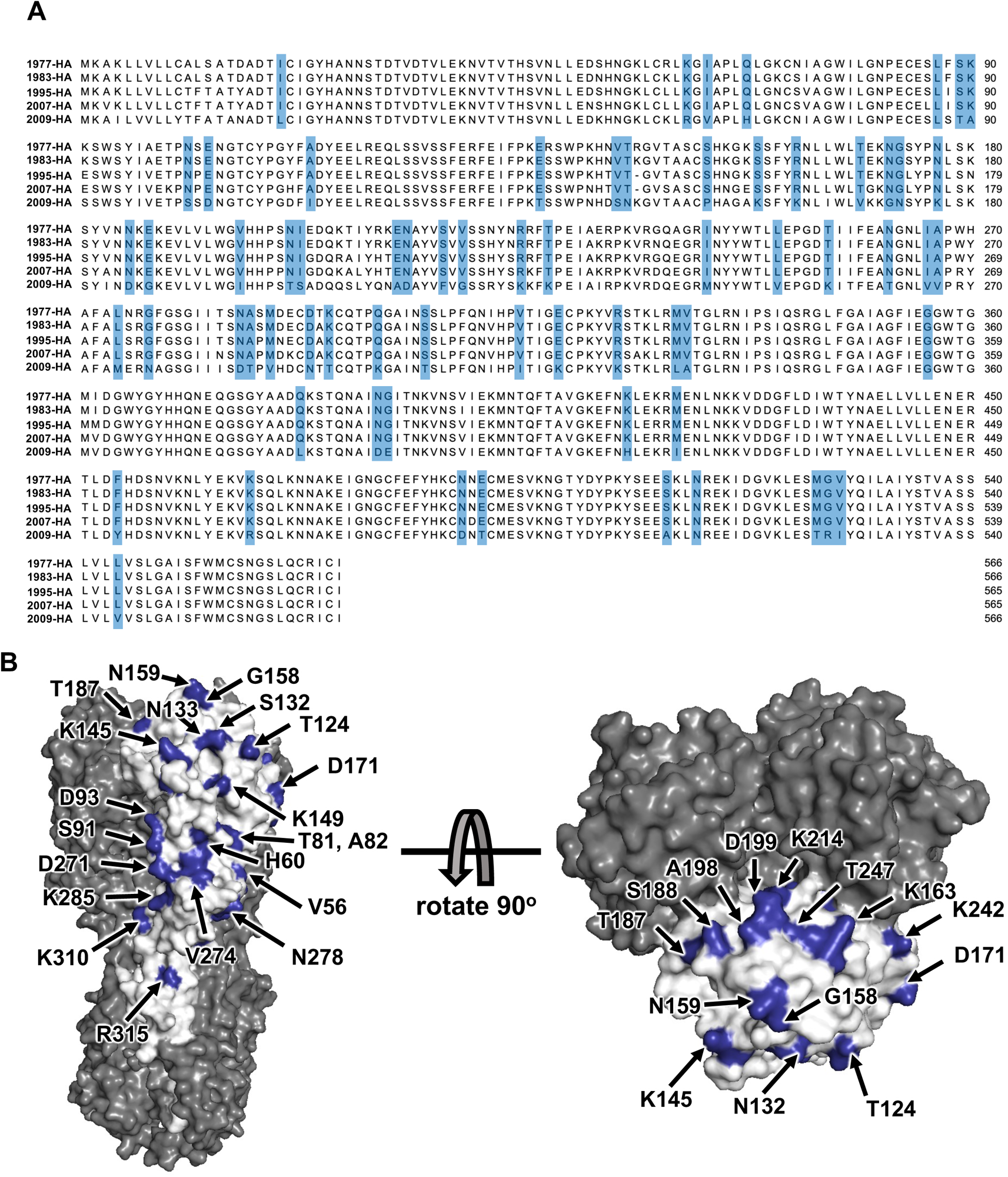
Surface residues difference among pre-2009 H1N1 HA and Cal/09 H1N1 HA. (A) Mutations are highlighted in blue on in a sequence alignment among four pre-2009 HA protein and Cal/09 HA. (B) Surface residues on Cal/09 HA head domain, which differs from four pre-2009 HAs, are highlighted on the Cal/09 HA protein.

